# The form and function of a multi-functional weapon system in male and female burying beetles (*Nicrophorus vespilloides*)

**DOI:** 10.1101/2025.11.21.689795

**Authors:** Jack M.L. Smith, Andrew M. Catherall-Ostler, Frederik Püffel, David Labonte, Rebecca M. Kilner

**Affiliations:** Department of Zoology, University of Cambridge, Downing Street, Cambridge, CB2 3EJ, United Kingdom; Department of Bioengineering, Imperial College London, London, SW7 2AZ, United Kingdom

**Keywords:** Animal weapon system, sexual dimorphism, contest behaviour, bite force, weapon-supportive trait, parental care, social evolution, burying beetle

## Abstract

Animal weapon systems are used for attack and defence during competition for resources including, though not confined to, competition for mates. They comprise the weapon itself and associated morphological structures – or ‘weapon-supportive traits’ – that are essential for the deployment of the weapon in combat. We investigate the form and function of a weapon system in burying beetles *Nicrophorus vespilloides*, to better understand why it differs between the sexes. Both males and females engage in contests with members of their own sex to monopolise a scarce carrion breeding resource. We show that mandibles (weapon during biting) and head width (weapon-supportive trait) are larger in males, and that males exhibit a disproportionately larger increase in bite force with head width than females. However, in staged contests with size-matched rivals of the same sex, the weapon system functioned in the same way for males and females: for each sex, the combined effects of head width and maximum bite force best predicted contest outcome. We suggest that the component parts of the weapon system each serve multiple additional functions, including tasks associated with parental care, which contribute differently to fitness in each sex. The resulting divergent selection pressures may explain why sexual dimorphism persists.

## INTRODUCTION

Animal contests are widespread in nature, and many animal taxa engage in fights with conspecifics over access to limited resources, especially when resource access is a major constraint on fitness [1]. Males of diverse species have evolved specialised weaponry to increase their success in fights over access to females [2–4]. Classic examples include exaggerated morphologies such as the branching antlers of male deer and the forked head-horns of male rhinoceros beetles [5–8]. However, males are not unique in their capacity to bear weapons. Female-female contests can also arise over key breeding resources such as territory, food and shelter, and can likewise impose selection on female morphology [1, 9, 10]. In some cases, females are better armed than males, such as females of the dung beetle *Onthophagus sagittarius* which have head-horns that are used in competition for breeding resources [11, 12]. Generally, though, female weapons are typically less developed than their male counterparts [8], and a key challenge is to explain why this sexual dimorphism in weaponry persists. At least three non-mutually exclusive explanations have been advanced to resolve this puzzle.

Since Darwin’s *Descent of Man* [13], the traditional explanation is that selection for exaggerated morphology is weaker in females because they less frequently engage in competition for mates; female weapons are thus less likely to contribute to fitness. For example, female reindeer have much smaller antlers than males, and do not use them in battles for access to mates, but instead in fights for social dominance and contend for scarce resources [8, 14].

A second argument relates to the more recent observation that weapons do not function in isolation but are instead integrated into a wider phenotype [15]. Thus, rather than focusing on the weapon *per se*, one ought to consider the wider ‘weapon system’ which includes the weapon-supportive traits that have evolved to enhance the use of the weapon itself [15]. Weapon-supportive traits include the anatomical and behavioural traits that support the effective deployment of a weapon during a contest and can thus, for example, be assessed through biomechanical measures related to force output [4, 16]. It may be that divergent selection pressures between the sexes constrain the evolution of the weapon-supportive traits, in turn influencing the size of the weapon that can be wielded by each sex.

A third, related consideration is that the different elements that make up a weapon system commonly serve more than just one function [15]. The particular array of functions served by each component of the weapon system could differ between the sexes, and so be the source of the constraints that limit the size of the weapon, or the weapon-supportive trait, or both.

Understanding the selective forces acting on the different structures in weapon systems is therefore key to understanding their relative size in each sex, and is also necessary to uncover the relationship between form and function. Fighting can involve actions such as grabbing, levering, shoving, striking, and piercing – all of which involve the targeted application of forces onto an opponent via a mechanical weapon. Mechanical weapon systems are specialised to deliver these forces. Investigating the underlying biomechanical function of the constituent parts is crucial to fully understand weapon system evolution and diversity [17].

Here, we describe experiments that relate the form of weapon system components to their function in the burying beetle *Nicrophorus vespilloides*. This species relies upon small vertebrate carrion to complete its lifecycle and it exhibits elaborate biparental care [18]. Both parents use their mandibles to strip the dead body of its feathers or fur, before rolling the body into a compact brood ball which is then coated with antimicrobial exudates [19]. The carcass is buried in a shallow crypt around which the female lays her eggs. Larvae hatch in the soil and crawl to the balled carcass, which becomes their edible nest and the site for parents to provide post-hatching care. Both parents commonly care together for larvae, provisioning them and defending them as they develop, though uniparental care is also seen [20, 21].

The burying beetle’s dependence on a scarce, unpredictable bonanza resource for breeding sets the stage for contests in both males and females. Contests frequently take place following carcass discovery, when rivals contest the right to monopolise the carrion breeding resource for their own reproduction. At this point, contests are typically staged within each sex, and the winners from each sex become the dominant pair on the dead body [20, 22]. Contests may also happen after larval hatching, when parents seek to protect their nest and larvae from infanticidal takeover by rival *Nicrophorus* conspecifics and heterospecifics [22]. Both sexes engage in resource defence, though males specialise in these duties of care when both parents are present [20, 23–25].

Previous work suggested that the burying beetle weapon system involves the mandibles as a primary mechanical weapon, because they inflict physical harm through biting during contests. Mandibles play an important role in combat across insect taxa [3, 8, 26, 27]. For example, in crickets, longer mandibles contribute to male fighting success [27–29]. Biting is also the main form of attack in burying beetles, causing cuticular damage to rivals, sometimes as extreme as decapitation, or in some cases, cannibalism, with the abdomen pierced and consumed in its entirety (JMLS pers. obs.).

The burying beetle mandible is wielded by weapon-supportive traits that often correlate with body size, and indeed, relative size is known to predict the outcome of contests within each sex [24, 30, 31]. In other insects, head size is often considered to be part of an individual’s weapon system, alongside biting and piercing mouthparts [26–29], because of its positive correlation with bite performance: larger heads likely host a larger volume of muscle to drive the mandibles, and thus result in increased bite performance [26, 32–36]. *N. vespilloides* exhibits sexual dimorphism in head width – a trait which serves as a reliable proxy for bite force in insects [37] – and this dimorphism is also found in several other species of *Nicrophorus* [38]. Although it has been hypothesised that a wider head evolved in males under selection from his greater engagement in contests [37], it remains unclear whether head width is indeed associated with fighting success.

In this study, we investigate the form and function of the weapon system in male and female burying beetles. Specifically (1) we investigate whether there exists sexual size dimorphism in the mandibles (the weapon), and how they scale with body size in each sex. We then (2) determine whether head width functions as a weapon-supportive trait, by quantifying the link between head width and biting performance in both males and females. Finally, we (3) assess the integrated function of the weapon system by staging experimental contests between sex-matched pairs. We investigate how linear measures of weapon size, i.e., mandible length, and the weapon support system, i.e., head width, jointly influence bite performance and can thus serve as predictors for contest outcome. By evaluating these traits together, we determine their relative contributions to contest success in each sex, leading to a better understanding of why sexual dimorphism persists in these traits.

## METHODS

### Experimental populations

The experiment used beetles from five wild populations and one laboratory stock population. Wild populations were collected from five localities at the intersection of Cambridgeshire, Bedfordshire, and Huntingdonshire (UK): Cockayne Hatley Wood (52°08′21″N, 000°09′04″W), Gamlingay Wood (52°09′54″N, 000°11′06″W), Hayley Wood (52°09′36″N, 000°06′53″W), Waresley Wood (52°10′33″N, 000°09′31″W), and Weaveley Wood (52°10′18″N, 000°12′49″W); they were bred for a single generation in the lab prior to the experiments. The large, genetically diverse stock population was 15 generations old at the time of our experiment and descended from crosses of wild beetles collected from three areas in Cambridgeshire/Huntingdonshire (UK): Gamlingay Wood (52°09′54″N, 000°11′06″W), Madingley Wood (52°13′02″N, 000°02′55″E), and Waresley Wood (52°10′33″N, 000°09′31″W). Prior to the experimental contests, adults were housed individually in plastic boxes (12 x 8 x 2 cm) and fed equal amounts of ground beef twice a week. In total, over 600 beetles were used in the experiments, with roughly half sourced from wild populations (males, N = 156; females, N = 168) and the other half from laboratory stock (males, N = 156; females, N = 168).

### Morphological measurements

Pronotum width was measured as a proxy for beetle body size, as is standard practice [39, 40]. Pronotum width was defined as the distance between the two widest points on the pronotum; head width was defined as the distance between the two widest points of the head capsule (described in detail in the study by Smith *et al.* [37]). Pronotum width, head width and mandible length were measured for all specimens using Mitutoyo digital callipers (accurate to 0.01 mm), before any bite force measures were taken or contests were staged.

Due to their small size, mandible measurements could only be taken destructively and so were made after experimental contests had ended. Beetles were frozen at -20 °C, and mandibles were dissected out under a stereomicroscope using entomological pins and fine forceps. Mandibles were mounted flat on their dorsal side in clear nail polish (approximately aligned with the dominant plane of mandible movement), photographed at 20x magnification, and then measured using ImageJ. Mandible length, defined as the distance between the mandibular hinge joint and the distal tip of the mandible in the dominant plane of mandible movement, served as measure for the mandible “outlever” [26, 27, 33, 34, 41]. Left and right mandible length did not differ significantly in length (paired t-test, t_198_ = 0.08, p = 0.93), and the right mandible was thus arbitrarily chosen for subsequent analysis.

### Force measurements and bite performance

Bite forces were measured using a bespoke force transducer, based on a pivoting beam and capacitive force sensor (see studies by Püffel *et al.* [34, 41, 42] for further details; data acquisition frequency 33 Hz, resolution 2 mN). The setup included two small bite plates (length 1 mm, thickness 0.15 mm) that were grasped by the distal tip of the mandibles; one plate is fixed in place, whereas the other is connected to a rigid beam, mounted onto a pivot. A bite pivots the beam such that it presses onto the capacitive force sensor (for a more detailed description of the experimental setup see studies by Püffel et al. [34, 41]). Because bite force depends on mandible opening angle [34, 43], the distance between the two bite plates was individualised for each beetle via a motor-controlled system, such that the gap was about 0.38 head width, chosen because it provided quality force readings across the size range. At comparable relative bite gaps (0.35-0.40), *Atta* ants bit with an average mandible opening angle of 80 ± 4° (n = 28, [41]), generating distal bite forces of 73 ± 14% compared to an extrapolated maximum force at smaller opening angles. The uniformity of the bite angle across beetle body sizes brought the further advantage of simplifying downstream analysis: both the angle between the mandible and the bite plate, and point of contact between mandible and plate were constant across trials, standardising the experimental conditions (for a more detailed discussion of these points, see Püffel et al. [41]).

To conduct bite force measurements, forceps were used to clasp beetles securely around the elytra with legs securely tucked behind, so that the head could be moved towards the bite plates unencumbered. When placed in front of the bite plates, there was variation in the latency to bite across individuals, perhaps reflecting differences in aggression. To account for this variation in downstream statistical analysis, bite latency was measured as the time between placing an individual in front of the sensor to the first bite; the first bite could be clearly identified by direct observation as it and movement of the bite plate were both clearly visible. Thereafter, bite forces were recorded for 30 seconds, after which the beetle was returned to its individual container. All individuals were measured once, as previous work in *Atta* ants revealed the effect of trial number on maximum recorded bite force [34].

The output of bite force measurements was a force trace for each beetle; a custom R script was used to extract three variables from each trace: (1) the latency to bite in seconds; (2) the maximum bite force in mN; (3) the bite rate as number of bites per second (calculated as the total number of bites divided by the duration of the interval between the first and last bite). A bite was defined as a peak in the force trace exceeding 30% of the maximum value measured during the trial.

### Experimental contests and predictors of contest success

Pairs of contestants were established within each wild population, or within the stock population. We used pronotum width to match contestants for body size [44] and ensured that any difference did not exceed one SD of the whole experimental sample (mean ± SD: 4.71 ± 0.37 mm). Consequently, there was no significant difference in the pronotum width between paired opponents (paired t-test, t_234_= 0.69, p = 0.49), and the magnitude of differences in pronotum width between opponents did not differ between populations (ANOVA, F_5_ = 1.07, p = 0.38). This design therefore successfully excluded any effects of differences in body size – a trait which is an important predictor of contest outcome in burying beetles [24, 30, 31] – and permits investigation of how head width, mandible length, and bite force influenced contest outcomes independent of body size differences. In total, 235 successful experimental contests were staged: 116 between males, and 119 between females.

The experimental contest protocol was informed by the work of Lee et al. [45]. Rival beetles were placed simultaneously in a plastic box (17 x 12 x 6 cm), previously filled with 2 cm of soil; a freshly thawed mouse was placed in the centre (mean mass = 24.6 ± 1.09 g). The box was then placed in a dark cupboard and left undisturbed for 72 h (approximately sufficient time for a contest to be concluded [24, 46–48]): if an individual was on the carcass whilst the other was far away in the soil, the former was designated the winner and the latter the loser. In the occasional instances when both beetles were found on the carcass, or when neither was present, the boxes were returned to the dark cupboard for an additional 24 h to allow more time for the contest to conclude. Any pairs in which there was still no clear winner after the additional 24 h were classed as a ‘no contest’ and excluded from the analysis. In total, 16 contests ended without resolution: 8 between males, and 8 between females.

### Statistical analysis

Analyses were performed using R version 4.4.0. We used ordinary least squares (OLS) and standardised major-axis (SMA) regression using the R package ‘*smatr*’ to analyse allometric relationships [49–51]. The suitability of these models is subject to debate, as they involve assumptions on ‘observational’ and ‘biological’ errors [49]. The text reports the results for OLS regressions for simplicity; SMA regression results are provided in the electronic supplementary material. Fortunately, the main conclusions of this paper are supported by either model; only a less central result depended on the regression model, and is thus described with both OLS and SMA regression. Allometry describes the relationship between a biological variable *y* (such as a morphological trait or performance measure) and body size *x*, typically assumed to follow a power law, *y* = *αx^β^*, where *α* is the allometric intercept, and *β* is the allometric scaling parameter. The relation between y and x is typically linearised through log-transformation, log(*y*) = log(*α*) + *β* log(*x*); this procedure enables determination of the scaling parameter via least-square regression that minimises the relative rather than the absolute error. Sex differences in biting behaviour were assessed using independent two-sample t-tests to compare latency to bite and bite rate between males and females.

Generalised linear models (GLM) with binomial error distribution and logit link function were used to identify the variables that best predicted contest outcome. Logistic regression analysis has historically been the preferred method for the analysis of animal contest outcomes [1, 45, 52]. To conduct this analysis, we first arbitrarily designated one individual from each fight as the ‘focal’ individual and the other as the ‘opponent’. Contest outcome, recorded as the dependent variable, was expressed in binary format: a value of 1 vs 0 indicated the focal individual vs the opponent won, respectively. For each contest, relative pairwise trait differences between the focal and opponent were calculated for head width, mandible length, maximum bite force, bite latency, and bite rate. These relative pairwise trait differences were then used as independent variables [1]. We calculated the relative difference by subtracting the opponent trait from the focal trait and then dividing this value by the mean of the two traits.

To determine the best predictors of contest success, we began with a global model that included all relative pairwise trait differences in head width, mandible length, maximum bite force, bite latency, and bite rate. We then performed model selection using the *dredge* function in the R package ‘*MuMIn*’ [53] to run a complete set of models with all possible combinations of predictors. Models were ranked according to the Akaike information criterion (AIC), and the model with the lowest AIC score was designated best in class [1, 54–56]. However, any model with an AIC with a difference of 2 or less from the best-fitting model was considered to be an equivalent model, and examined in more detail [4, 57]. Next, to avoid retaining unnecessarily complex models, we implemented a nesting rule, removing any model that was a more complex version of a simpler model with a lower AIC value [57–59].

Variance inflation factors (VIFs) were calculated using the *check_collinearity* function in the R package ‘*performance*’ [60], which assesses multicollinearity among predictors. A VIF of 1 indicates no collinearity, and a VIF > 10 suggests strong collinearity [52]. In all cases, the VIF values were low, ranging from 1.02 to 1.31.

## RESULTS

### (1) Sex differences in the scaling of head traits

#### a) Mandible length

For the same body size, males consistently displayed longer mandibles than females, regardless of the type of regression analysis used. However, the scaling relationship between body size and mandible length differed with regression analysis.

Using OLS, mandible length exhibited a negative allometric relationship with pronotum width in both males (OLS slope = 0.88, 95% CI [0.80, 0.97], F_242_ = 7.10, p < 0.01) and females (OLS slope = 0.88, 95% CI [0.79, 0.96], F_252_ = 7.78, p < 0.01). Although there were no significant differences in slopes between the sexes (OLS: t_494_ = -0.09, p = 0.93), the elevation of the slope was significantly greater for males than for females (OLS: t_495_ = -2.81, p < 0.01).

However, using SMA, mandible length exhibited a positive allometric relationship with pronotum width in both males (SMA slope = 1.12, 95% CI [1.03, 1.21], r_242_ = 0.18, p < 0.01, test against isometry [slope = 1]) and females (SMA slope = 1.12, 95% CI [1.04, 1.21], r_252_ = 0.18, p < 0.01, test against isometry [slope = 1]). Once again, there were no significant differences in slopes between sexes (SMA: LR_1_ = 0.01, p = 0.94; Figure 1a), and the elevation of the slope was significantly greater for males than for females (SMA: χ^2^_1_ = 6.82, p < 0.01).

**Figure 1.**
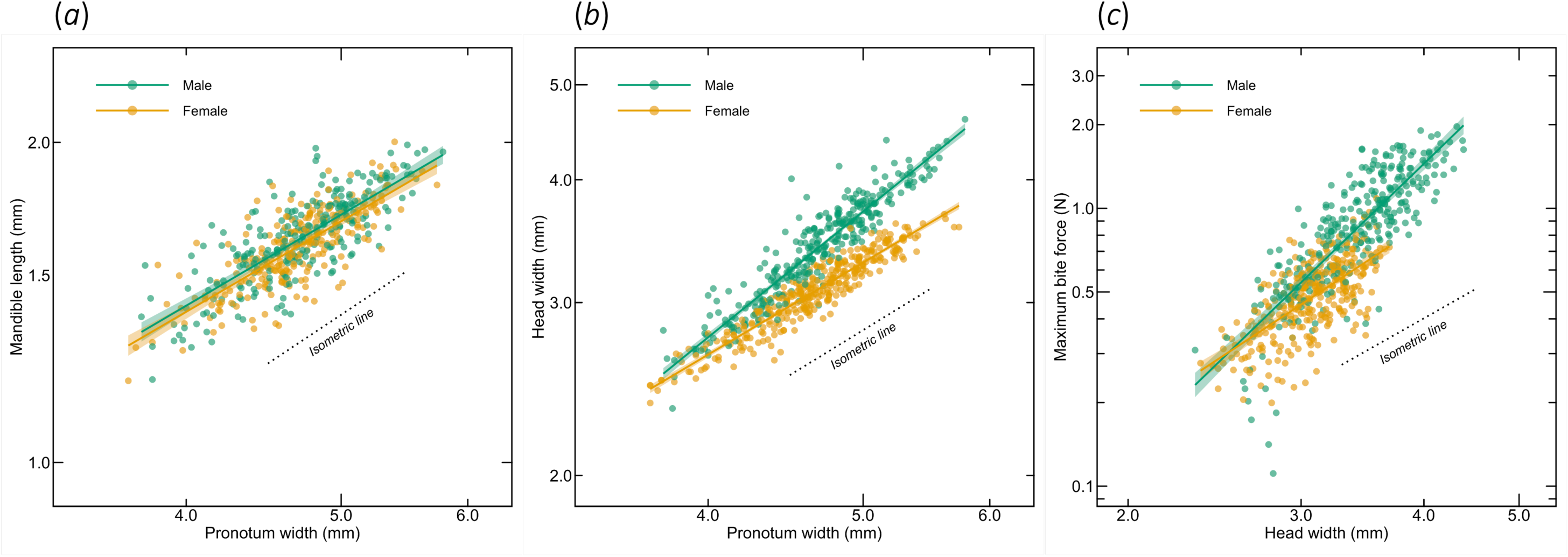
Allometric relationships in male (green) and female (orange) *Nicrophorus vespilloides.* Lines shown are the results of ordinary least-squares regressions on log–log transformed data with 95% confidence intervals, dotted lines show predictions from isometry. **(a)** The allometric relationship between mandible length and pronotum width. Mandible length scaled with negative allometry in both sexes, with similar slopes but a significantly higher elevation in males. **(b)** The allometric relationship between head width and pronotum width. Head width scaled with positive allometry in males but exhibited an isometric relationship in females. **(c)** The allometric relationship between maximum bite force and head width. Bite force scaled with strong positive allometry in both sexes, but the slope was significantly steeper in males.

#### **b)** Head width

In line with previous work by Smith et al. [37], we found that head width scaled isometrically with body size in females, and that the relationship was significantly steeper in males. Head width exhibited a positive allometric relationship with pronotum width in males (OLS slope = 1.32, 95% CI [1.27, 1.38], F_310_ = 130.38, p < 0.001) and an isometric relationship in females (OLS slope = 0.97, 95% CI [0.93, 1.01], F_334_ = 2.86, p = 0.09). The slopes consequently differed significantly between the sexes (OLS: t_644_ = -10.51, p < 0.001; Figure 1b).

### (2) Sex differences in the relationship between head width and biting performance

#### a) Bite force

Individuals with wider heads exerted a greater bite force. This increase was larger in males than in females. Bite force increased significantly with head width in males (OLS slope = 3.42, 95% CI [3.16, 3.68], F_308_ = 117.22, p < 0.001) and females (OLS slope = 2.33, 95% CI [2.05, 2.60], F_334_ = 5.45, p = 0.02). For both sexes, bite forces were positively allometric (the isometric expectation is a scaling coefficient of 2), but the extent of this positive allometry was significantly stronger in males (OLS: t_642_ = -5.56, p < 0.001; Figure 1c).

#### **b)** Latency to bite and bite rate

There was a significant difference in the latency to bite between the sexes (t-test, t_622.25_ = 2.12, p = 0.03). The male latency to bite was 7.99 ± 0.69 seconds on average, about 30% slower than the average female latency (6.07 ± 0.59 seconds). Bite rate, however, did not differ significantly between the sexes (t-test, t_454.31_ = -1.57, p = 0.12; 0.56 ± 0.03 vs 0.60 ± 0.01 for males and females, respectively).

### (3) The predictors of contest outcomes

A model that included both head width and maximum bite force as explanatory variables provided the best predictions of contest outcome, as indicated by the lowest AIC score (Table 1). Both head width and maximum bite force were significant predictors of contest outcome when included in the same model (relative head width difference: z_232_ = 5.00, p < 0.001; relative maximum bite force difference: z_232_ = 2.69, p = 0.01; Figures 2 and 3). The magnitude of these effects can be illustrated by extracting the probability statistics from the logistic regression (Tables 2 and 3). For contests where individuals were equally matched in head width and bite force (i.e., a relative difference of 0), the predicted probability of winning a contest was 0.48 (95% CI [0.41, 0.55]; Tables 2 and 3) – random, as expected. A difference of 15% in head width was sufficient to all but guarantee a victory (91% chance of winning, 95% CI [0.80, 0.96]). To achieve a comparable success chance via a difference in bite force required a much larger 150% increase (95% CI [0.62, 0.98]).

**Table 1.** Results of generalised linear models used to investigate which relative trait differences best explain contest success in *Nicrophorus vespilloides*. Models were ranked by AIC, and only models within two units of the best-fitting model (!AIC ≤ 2) are presented, including the null (intercept only) model for comparison. Relative differences were calculated by subtracting the opponent trait from the focal trait and then dividing this value by the mean of the two trait values. The best-supported model incorporating head width and maximum bite force provided the best prediction of contest outcome, outperforming all other variables or combinations of variables. HD, head size difference; FD, maximum force difference; BRD, bite rate difference; MD, mandible length difference; BLD, latency to bite difference; k, number of parameters; LL, log likelihood; AIC, Akaike information criterion, ΔAIC, distance of each candidate model from best fit model as indicated by AIC scores; w, Akaike weight.

| Model | k | LL | AIC | $\Delta\text{AIC}$ | w |
| --- | --- | --- | --- | --- | --- |
| Outcome ~ HD + FD | 3 | -128.26 | 262.51 | 0 | 0.30 |
| Outcome ~ HD + FD + MD | 4 | -127.98 | 263.96 | 1.45 | 0.15 |
| Outcome ~ HD + FD + BLD | 4 | -128.03 | 264.06 | 1.55 | 0.14 |
| Outcome ~ HD + FD + BRD | 4 | -128.03 | 264.06 | 1.55 | 0.14 |
| Null | 1 | -162.87 | 327.74 | 65.23 | 0.00 |

**Figure 2.**
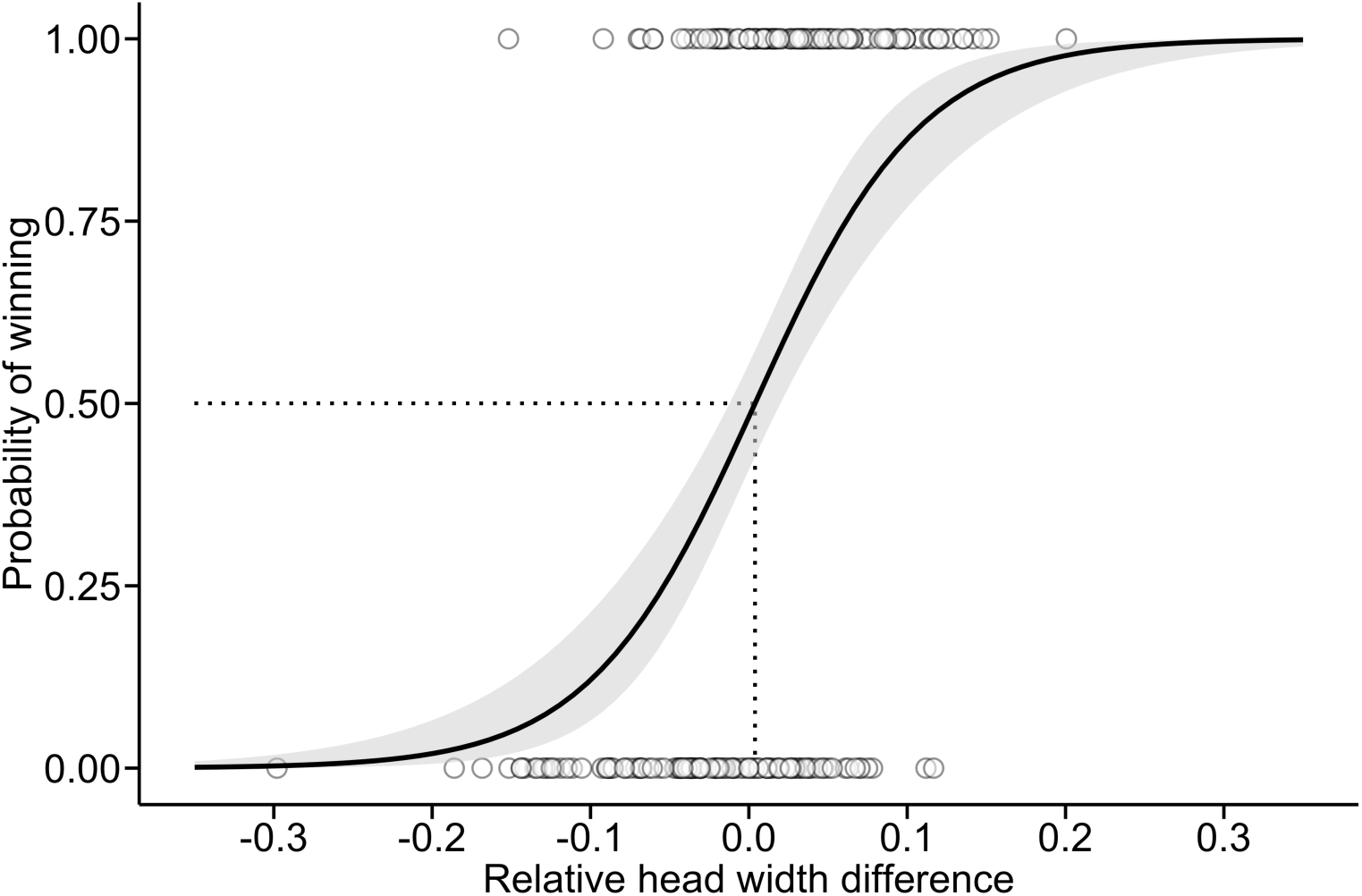
Logistic regression showing the relationship between an individual’s probability of winning a contest and head width in *Nicrophorus vespilloides* (sexes pooled). Shaded region shows 95% confidence intervals, dotted lines indicate the inflection point of the logistic regression i.e., the relative head width difference where the probability of winning a contest is 50%.

**Figure 3.**
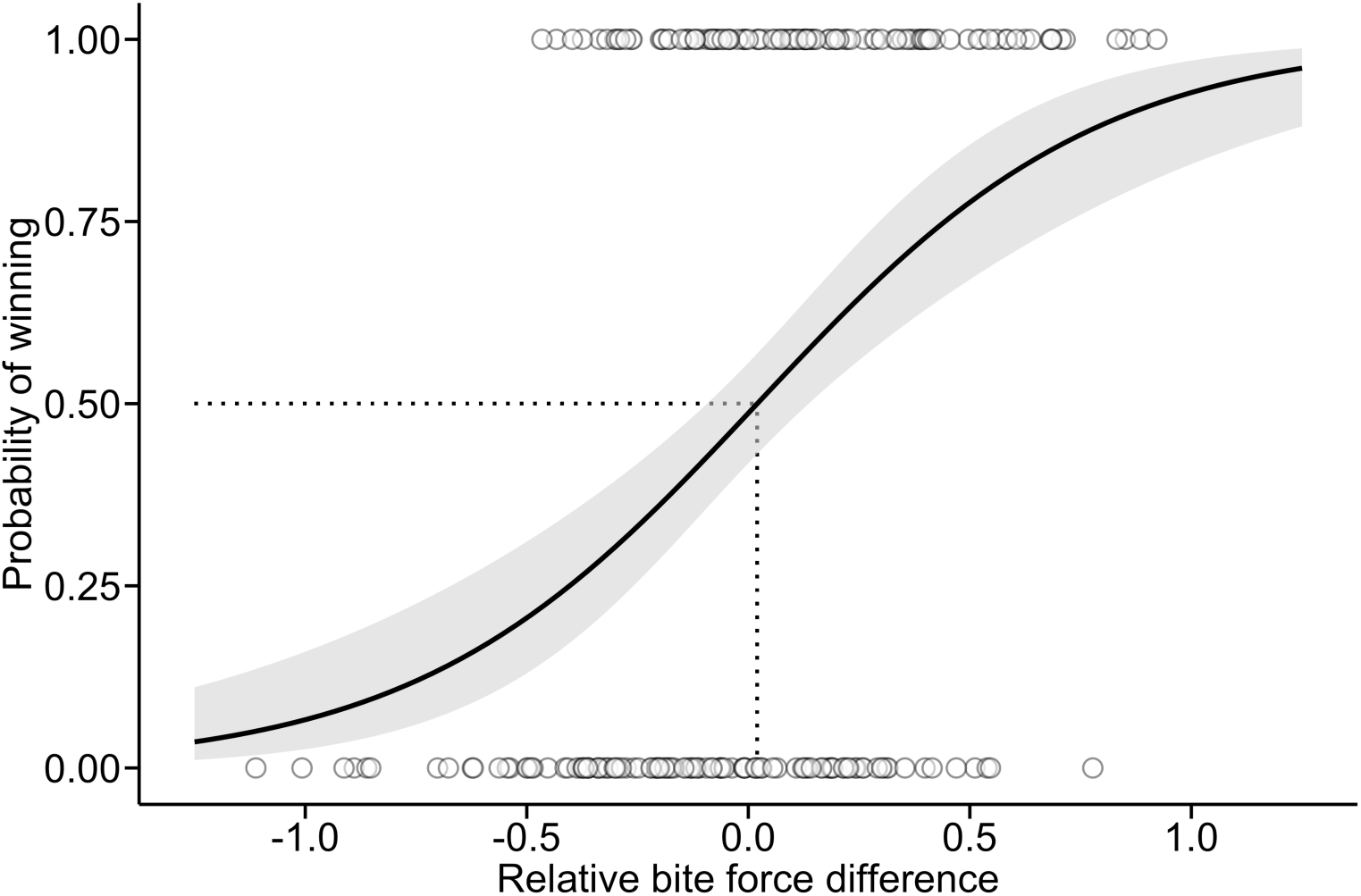
Logistic regression showing the relationship between an individual’s probability of winning a contest and maximum bite force in *Nicrophorus vespilloides* (sexes pooled). Shaded region shows 95% confidence intervals, dotted lines indicate the inflection point of the logistic regression i.e., the relative bite force difference where the probability of winning a contest is 50%.

**Table 2.** Predicted probabilities showing how a relative increase in head width influences the probability of winning a contest. Predictions were generated using the best-fitting logistic regression model, with relative bite force difference held constant at zero. At a head width difference of 0.00, the predicted probability of winning a contest was 0.48, 95% CI [0.41, 0.55], indicating that when opponents were perfectly matched in head width, both had an approximately equal probability of winning.

| Relative head width difference | Probability of winning | 95% CI |
| --- | --- | --- |
| 0.00 | 0.48 | 0.41, 0.55 |
| 0.02 | 0.56 | 0.48, 0.64 |
| 0.05 | 0.67 | 0.58, 0.76 |
| 0.10 | 0.82 | 0.70, 0.90 |
| 0.15 | 0.91 | 0.80, 0.96 |
| 0.20 | 0.96 | 0.87, 0.99 |
| 0.30 | 0.99 | 0.95, 1.00 |

**Table 3.** Predicted probabilities showing how a relative increase in bite force influences the probability of winning a contest. Predictions were generated using the best-fitting logistic regression model, with relative head width difference held constant at zero. At a bite force difference of 0.00, the predicted probability of winning a contest was 0.48, 95% CI [0.41, 0.55], indicating that when opponents were perfectly matched in bite force, both had an approximately equal probability of winning.

| Increase in relative bite force | Probability of winning | 95% CI |
| --- | --- | --- |
| 0.00 | 0.48 | 0.41, 0.55 |
| 0.20 | 0.55 | 0.46, 0.64 |
| 0.50 | 0.66 | 0.51, 0.78 |
| 1.00 | 0.80 | 0.57, 0.92 |
| 1.50 | 0.89 | 0.62, 0.98 |
| 2.00 | 0.95 | 0.67, 0.99 |
| 2.50 | 0.97 | 0.71, 1.00 |
| 3.00 | 0.99 | 0.75, 1.00 |

Although several other models had similar AIC values (ΔAIC > 2; Table 1), all were more complex versions of the top model, and none of the added predictors reduced AIC enough to clearly justify the increase in model complexity, in accordance with the aforementioned nesting rule (see *Methods*; [57–59]).

To investigate potential sex differences in these predictors, we added interaction terms to the candidate model. However, neither interaction was statistically significant (head width difference*sex: z_229_ = -1.61, p = 0.11; maximum bite force difference*sex: z_229_ = -0.70, p = 0.48), and the model with interactions had a higher AIC compared to the candidate model without interactions. We conclude that both head width and maximum bite force contribute to contest success in a similar way for each sex.

## DISCUSSION

Understanding the evolution of animal weapon systems involves relating the form of the weapon system to its function and evaluating how it contributes to fitness within a behavioural and ecological context [15]. In insects, relatively few studies have explored how the distinct components of the weapon system relate to biomechanical performance, and how they therefore contribute to success in a contest. Rarer still are studies which investigate these relationships in both sexes. Addressing both shortcomings, we here showed how different components of a weapon system function in each sex, in relation to their form, and how they combine to influence the outcome of male-male and female-female contests in the burying beetle *N. vespilloides*.

### Sex differences in the scaling of head traits

We found that bite force was a key predictor of success in a contest, thus confirming that the mandibles are the mechanical weapons at the centre of the weapon system we analysed here, and in common with many other insect taxa in many insect taxa [3, 8, 26, 27]. We found that the mandibles themselves scaled close to isometry with body size, though were consistently smaller in females. Perhaps surprisingly, their relative size did not predict the likelihood of winning a contest when other components of the weapon system were used as covariates in the regression. One explanation could be that the mandibles serve multiple purposes, since they are used in nest maintenance and carcass preparation, as well as in contests. This could explain why the mandibles are larger in males than in females, given the male’s greater effort in carcass preparation [61], though the intricate process of preparing a brood ball and stripping it of its fur or feathers could be hindered by enlarged mandibles, and potentially place an upper limit male mandible length. It is notable that insect species bearing greatly enlarged mandibles, such as many Lucanidae (e.g., *Lucanus cervus*) undertake combat in open spaces where males can use their elongated mandibles to maintain distance, or to signal their fighting prowess before any battle commences [15]. In contrast, burying beetles inhabit underground crypts where contests over nest and brood defence occur at close quarters [20], which might select against both the deployment of larger mandibles in a contest and their use as a visual display of fighting ability.

Further differences between the sexes were observed in a key weapon supportive trait analysed here: head width. Consistent with patterns previously described in different populations of *N. vespilloides* [37], male head width exhibited positive allometry, whereas in females, head width scaled isometrically. One possible explanation for this divergence lies in the balance of selective pressures shaping head width in the two sexes. Perhaps there are divergent selection pressures on the sexes that are linked to task specialisation during biparental care [62–64]. For example, before becoming primarily responsible for nest and brood defence, males take the lead in transforming carrion into an edible nest for their larvae, a task that involves chewing off fur or feathers as well as powerful bites to shape the carcass into a compact and rounded ball [20, 65, 66]. Larger males, presumably with correspondingly larger heads, are more effective at preparing the carrion ball [61, 67]. The combined demands of preparing the carrion nest and then defending it may have selected wider heads in males, since a greater bite force would be advantageous in both contexts.

By contrast, the isometric scaling of female head width could suggest a balance between different selective pressures [68] that select for or against positive allometry [69, 70]. Females play a greater role in offspring provisioning and post-hatching carrion maintenance [66, 71]. A narrower head in females may enhance their ability to provide care to offspring, such as transferring fluids to larvae though oral trophallaxis, perhaps by improving the alignment of mouthparts between mother and offspring, thereby potentially stimulating a stronger feeding response and ensuring more efficient fluid transfer [37]. The costs of manufacturing a larger head could also differ between the sexes. Perhaps the costs of expenditure on weapon systems by females are disproportionately greater than in males, depressing both female fecundity and parental investment [9], much as fecundity trade-offs in females constrain the evolution of sexually selected traits in other species [72, 73]. Female weapons of snapping shrimp *Alpheus heterochaelis* exhibit negative allometry, for example, because snapping shrimp reduce relative claw size during the breeding season to increase investment in egg production and brood care [74]. Further work is required to test whether the dimorphism observed in head width is indeed related to task specialisation in each sex, and the different functions that this trait supports.

### Sex differences in the relationship between head width and biting performance

Studies on other insects have found that head size is an important component of insect weapon systems through its influence on bite force [26, 27, 29]. Head width is thought to be a good predictor of bite force, because it is proportional to the assumed volume of muscle within the head capsule [26, 33–36] and thus the muscle cross-sectional area, which is a more direct predictor of bite force [33, 34]. It is possible that other morphological attributes are linked to the muscles within the head and could additionally contribute to bite force, such as head height or length. Additional factors that could contribute to bite force, but which might not be detected with head measurements, include changes in the arrangement of the head muscles (e.g., shorter muscles with larger cross-sectional areas) or an increased mechanical advantage (ratio between mandible inlever and outlever [34]).

The relationship between head width and maximum bite force differed between the sexes. Although head width predicted biting maximum force in both males and females, each incremental increase in head width yielded a larger incremental increase in maximum bite force in males than in females. Regression lines for the two sexes intersect at the lower end of the head width range, indicating that small males and females of equivalent head width produce similar bite forces. The male advantage in bite force emerges only among larger headed individuals, suggesting that sex-specific specialisation for high bite forces only relates to the upper size range, where males seem to invest more heavily than females. It is possible that the mandible closer muscle is larger in males than females of an equivalent size, that males have a different muscle ultrastructure with longer sarcomeres [32, 34, 75, 76], or that that males have more densely packed muscle fibres [26]. Perhaps males are more strongly selected to have a high bite force because they invest relatively more effort in defending the brood and nest from rivals and therefore stand to gain more fitness through these actions than do females [20, 23–25].

### The predictors of contest outcomes

We found that the combined effects of head width (weapon-supportive trait) and maximum bite force (weapon system function) provided the best prediction of contest outcome in both males and females (cf. [77]). As discussed above, it is possible that weapon size in other species is primarily used to signal fighting prowess, to settle disputes by assessment, avoiding escalation to a physical contest [4, 78]. Agonistic signalling is used by *N. americanus* during contests, for example. In this species, both male and female beetles bear extended clypeal membranes, located above their mandibles, to provide exaggerated visual cues of body size to rivals [79]. However, *N. vespilloides* does not possess any equivalent traits and therefore may place less emphasis on pre-fight signalling to resolve contests. Head width and bite performance might both be critical in determining the outcome of contests that are resolved exclusively through direct combat. This may have been more likely in our experimental contests since we chose rivals that were matched in body size [80].

Our results suggest that head width has additional functions in direct combat which are not solely related to bite force. In burying beetles, fighting includes not only biting but also pushing, and flipping opponents over [40]. The head may support such combative behaviours, as is the case in Tephritid fruit flies, where males lunge at and head butt opponents [81], or in field crickets, which, alongside, biting, kicking, grappling, also utilise head butting in their behavioural repertoire [82]. Head butting has also been observed in females of other beetle species, such as female rhinoceros beetles, which lack the extravagant horns of males yet still fight over access to feeding sites by striking each other with their heads [83]. Thus, it is possible that the head itself is a mechanical weapon in addition to a weapon-supportive trait – a hypothesis best pursued in further work.

Interestingly, the effects of head width and bite force on contest success were consistent between the sexes, suggesting that the mechanisms underlying contest resolution are shared between males and females, despite distinct differences in allometries. It is consistent with the suggestions that sexual dimorphism in head width may be driven by divergent task specialisation during biparental care, and perhaps independent of the contests that take place around initial carcass discovery.

In summary, we have investigated the form and function of the same weapon system in males and females. We have found that the component parts, i.e., the weapon (mandibles) and the weapon supportive trait (i.e., head width), are each larger in males than in females. However, the component parts of the weapon system function together in the same way for males and females in the way they contribute to the outcome of a contest. Sexual dimorphism could be attributable to the fact that each component of the weapon system serves multiple functions, not all related to fighting, with each function contributing differently to fitness in each sex. It is also possible that the head itself is a mechanical weapon, or that the head supports multiple weapons, though to different degrees in each sex.

Our findings are consistent with the hypothesis that task specialisation during biparental care imposes divergent selection on male and female head width. The challenge for future work is to test this interpretation more robustly, for example through long-term experimental evolution in which the division of parental duties between the sexes is manipulated across multiple generations, and to determine whether it applies to other species too.

## Acknowledgements

We thank the BCN Wildlife Trust for access to Gamlingay, Hayley and Waresley Woods; the Tetworth Estate for access to Weaveley Wood; and Mr Michael Astor for access to Cockayne Hatley Woods.

## Authors’ Contributions

Jack M.L Smith, Andrew M. Catherall-Ostler and Rebecca M. Kilner conceived the ideas and designed the methodology; Jack M.L Smith and Andrew M. Catherall-Ostler collected the data; Frederik Püffel and David Labonte designed and built the bespoke force transducer, and provided biomechanical advice on the experimental setup and data analysis; Jack M.L Smith analysed the data; Jack M.L Smith and Rebecca M. Kilner led the writing of the manuscript. All authors contributed critically to the drafts, review, and editing, and gave final approval for publication.

