## Supplementary material for "The form and function of a multi-functional weapon system in male and female burying beetles (*Nicrophorus vespilloides*)"

### Sex differences in the scaling of head traits (SMA results)

#### Head width

Head width exhibited a positive allometric relationship with pronotum width in males (SMA slope = 1.41, 95% CI [1.36, 1.47], $r_{310} = 0.71$, $p < 0.001$, test against isometry [slope = 1]) and an isometric relationship in females (SMA slope = 1.03, 95% CI [0.99, 1.07], $r_{334} = 0.08$, $p = 0.12$, test against isometry [slope = 1]). The slopes consequently differed significantly between the sexes (SMA: $LR_1 = 121.87$, $p < 0.001$).

#### Bite force

Bite force increased significantly with head width in both males (SMA slope = 4.12, 95% CI [3.87, 4.39], $r_{308} = 0.82$, $p < 0.001$) and females (SMA slope = 3.46, 95% CI [3.20, 3.75], $r_{334} = 0.61$, $p < 0.001$. Test against isometry for both [slope = 2]). For both sexes, bite forces were positively allometric, but the extent of this positive allometry was significantly stronger in males (SMA: $LR_1 = 11.41$, $p < 0.001$).
